# In vivo magnetic recording of neuronal activity

**DOI:** 10.1101/092569

**Authors:** Laure Caruso, Thomas Wunderle, Christopher Murphy Lewis, Joao Valadeiro, Vincent Trauchessec, Josué Trejo Rosillo, José Pedro Amaral, Jianguang Ni, Patrick Jendritza, Claude Fermon, Susana Cardoso, Paulo Peixeiro Freitas, Pascal Fries, Myriam Pannetier-Lecoeur

**Affiliations:** SPEC, CEA, CNRS, Université Paris-Saclay, CEA Saclay 91191 Gif-sur-Yvette Cedex, France.; Ernst Strüngmann Institute (ESI) for Neuroscience in Cooperation with Max Planck Society, Deutschordenstraße 46, 60528 Frankfurt, Germany.; Instituto de Engenharia de Sistemas de Computadores-Microsystems and Nanotechnology (INESCMN), Rua Alves Redol, No. 9, Lisboa 1000-029, Portugal.; Instituto Superior Técnico IST, Physics Department, Universidade de Lisboa, Lisbon 1049-001, Portugal.; Donders Institute for Brain, Cognition and Behaviour, Kapittelweg 29, 6525 EN Nijmegen, Netherlands.

**Author notes:** These authors contributed equally (Pascal Fries, Myriam Pannetier-Lecoeur).

**Keywords:** Magnetic fields, magnetoencephalography, MEG, spin electronics, magnetic sensors

## Abstract

Neuronal activity generates ionic flows and thereby both magnetic fields and electric potential differences, i.e. voltages. Voltage measurements are widely used, but suffer from isolating and smearing properties of tissue between source and sensor, are blind to ionic flow direction, and reflect the difference between two electrodes, complicating interpretation. Magnetic field measurements could overcome these limitations, but have been essentially limited to magnetoencephalography (MEG), using centimeter-sized, helium-cooled extracranial sensors. Here, we report on in vivo magnetic recordings of neuronal activity from visual cortex of cats with *magnetrodes*, specially developed needle-shaped probes carrying micron-sized, non-cooled magnetic sensors based on spin electronics. Event-related magnetic fields inside the neuropil were on the order of several nanoteslas, informing MEG source models and efforts for magnetic field measurements through MRI. Though the signal-to-noise ratio is still inferior to electrophysiology, this proof of concept demonstrates the potential to exploit the fundamental advantages of magnetophysiology.

**HIGHLIGHTS:**

- Spin-electronics based probes achieve local magnetic recordings inside the neuropil
- Magnetic field recordings were performed in vivo, in anesthetized cat visual cortex
- Event-related fields (ERFs) to visual stimuli were up to several nanoteslas in size
- ERFs could be detected after averaging less than 20 trials

**IN BRIEF:** Caruso et al. report in vivo, intra-cortical recordings of magnetic fields that reflect neuronal activity, using magnetrodes, i.e. micron size magnetic sensors based on spin electronics.

## INTRODUCTION

Neuronal activity entails ionic flows across the cell membrane and along dendrites. This electrical activity can be measured extra-cellularly or intra-cellularly by microelectrodes (Kandel et al., 2000) which are either thin metallic micro-wires, or glass pipettes containing an ionic solution, to realize a conductive interface between the local brain tissue and the recording instrumentation. Intracellular recordings directly reveal the transmembrane voltage or current of an isolated neuron, but intracellular recordings in vivo are difficult in practice and often only brief measurements of single neurons are feasible. Extracellular recordings, on the other hand, measure the aggregate fluctuations in voltage arising from the net neuronal activity around the electrode’s tip, with respect to a reference electrode (Buzsáki et al., 2012). Microelectrodes inside the neuropil record action potentials and local field potentials (LFPs), electrocorticographic electrodes provide mesoscopic LFPs, and scalp electrodes deliver the electroencephalographic (EEG) signal. Combining many electrodes into planar (Maynard et al., 1997) or laminar arrays (Lewis et al., 2015) allows for the study of whole brain networks and their dynamics in the intact brain (Buzsáki, 2004).

The electric currents flowing through the active neuropil also give rise to a magnetic signature. Magnetoencephalography (MEG) (Cohen, 1968, 1972; Hari and Salmelin, 2012) is a non-invasive method to measure the magnetic fields of active neuronal populations during perceptual or cognitive tasks in the healthy or diseased human brain. This technique uses Superconducting Quantum Interference Devices (SQUIDs) cooled down to the temperature of liquid helium (4.2 K). The apparatus necessary for this cooling imposes a distance to the cortical surface of 3 to 5 cm in in vivo configurations. The spatial resolution is typically better than for EEG recordings, but even under optimal conditions still lies in the order of several mm^3^, with signal amplitudes in the femtotesla (10^−15^T) to picotesla (10^−12^T) range.

Local magnetic recordings of the neuronal activity could be a complementary technique to electrophysiology, because the magnetic signal provides interesting properties in addition to those realized by the electric signal. Contrary to electric fields, which strongly depend on the dielectric properties of the tissue between neuronal sources and the recording electrode, magnetic fields travel through tissue without distortion, because the respective permeability is essentially the same as free space (Barnes and Greenebaum, 2007). Therefore, magnetic fields are only attenuated by the distance to the current source. Ionic flows and the corresponding magnetic fields are likely largest inside neurons. As those magnetic fields pass through the cell membrane without attenuation, extracellular magnetic field measurements might provide functionally intracellular measurements without impaling the neuron. Moreover, while electrophysiological recordings yield scalar values, local magnetic recordings yield information about both amplitude and direction of current sources. Thereby, they might allow the precise localization of the source of neuronal activity at a given moment in time in the 3D volume of the brain. Furthermore, electrodes always measure the electric potential relative to a reference electrode, and the position and type of reference can substantially influence the measured signal. Moreover, in multi-electrode recordings, all channels typically share the same reference, which poses a problem for analyses of functional connectivity, because the resulting signals are not independent. Magnetrodes, presented in this work, provide an elegant solution, because the recorded magnetic signals are reference-free, and therefore allow for an unbiased measure of connectivity and information flow throughout the brain. In addition, these magnetrodes can be used to perform magnetic resonance spectroscopy (Guitard et al., 2016).

In order to minimize tissue damages, magnetic probes for insertion into the brain require a needle shape and the miniaturization of the magnetic sensors, while maintaining a very high sensitivity at physiological temperature. Approaches to record the magnetic biological signal closer to the sources than MEG have been successfully realized by using small SQUIDs (Magnelind, 2006), atomic magnetometers (Sander et al., 2012) or winded coils (Roth and Wikswo, 1985) and very recently with nitrogen-vacancy centers in diamond on a living invertebrate (Barry et al., 2016). However, limitations due to the millimeter size of the sensors or to its operating conditions never allowed penetration into the neuropil nor recording at distances of merely tens of microns from active cells.

## RESULTS

### Development and fabrication of micron-size magnetic sensors based on spin electronics for in vivo recordings

Spin electronics (Baibich et al., 1988) offers the capability to reduce magnetic sensors to micron size and to reach sensitivity in the sub-nanotesla range while working at body temperature and thereby avoiding bulky vacuum isolation (Reig, 2013). We have designed Spin Valve (Dieny et al., 1991) Giant Magneto-Resistance (GMR) sensors consisting of 5 segments of 4x30 μm^2^ arranged in a meandering configuration on silicon substrate that was ground to a thickness of 200 μm and etched to form a needle shape for tissue penetration (Fig. 1A). The sensors have been electrically insulated by a dielectric bilayer of Si_3_N_4_/Al_2_O_3_. We refer to these probes as ‘magnetrodes’, for a magnetic equivalent of electrodes (see STAR Methods for details of manufacturing and characterization).

**Figure 1.**
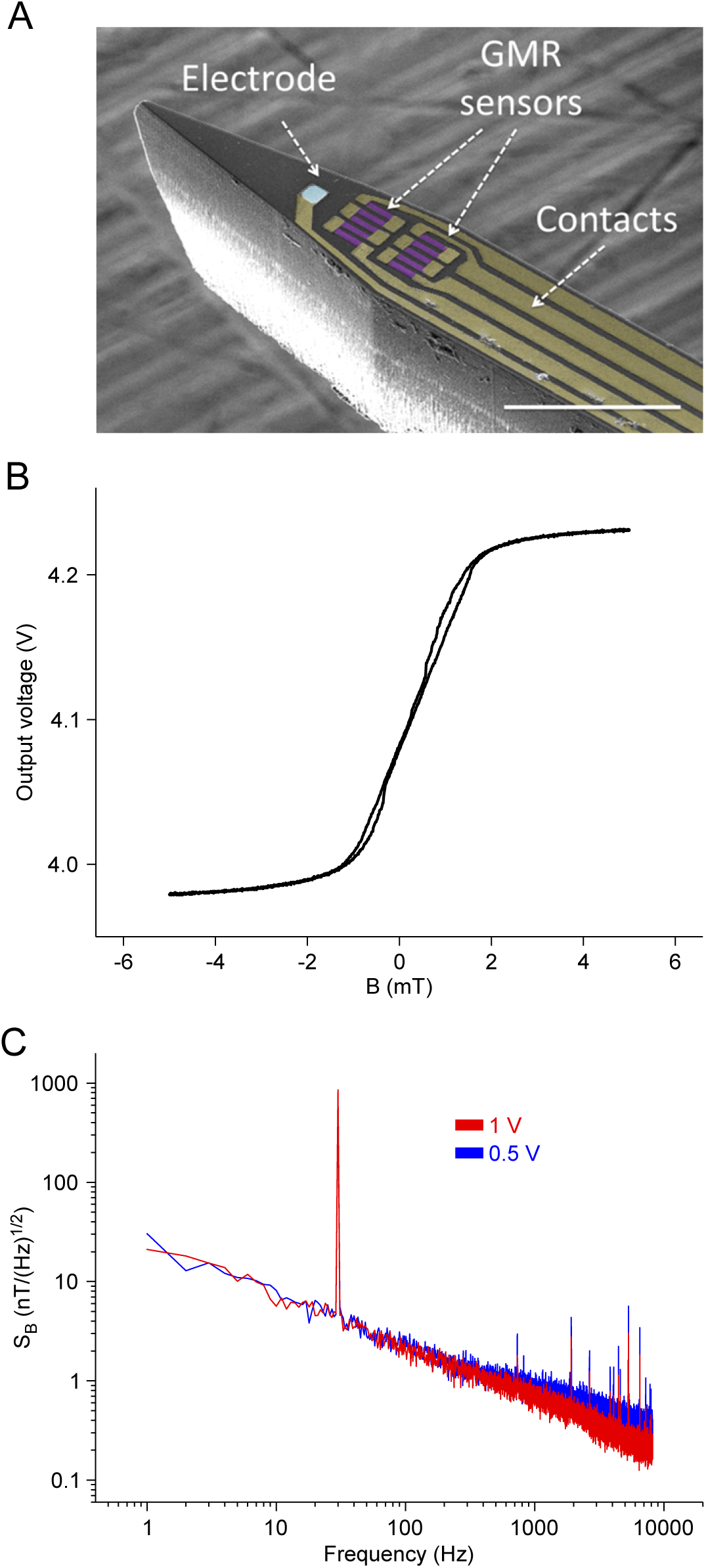
Magnetrode description and magnetic characteristics. (A) Scanning Electron Microscopy picture of a magnetrode containing 2 GMR elements, each with a meandering configuration. The elements are deposited on a 200 μm substrate that is 150 μm wide before narrowing at an 18° angle towards the tip. The sensitive direction is in the plane of the elements and orthogonal to the long axis of the tip. A platinum electrode (blue square) has additionally been deposited, but no recordings were achieved with it. Scale bar 100 μm. (B) Output voltage of the GMR sensor as a function of the magnetic field. The sensor is used for very weak magnetic fields around zero, which lead to outputs within the steep linear part of the curve. In the linear part, the slope is 1.8%/mT. (C) Equivalent-field noise spectral density S_B_ from 1 Hz to 10 kHz of the corresponding probe for 500 mV and 1 V peak-to-peak AC voltage of the GMR element. To obtain SB, the output voltage is converted in field-equivalent by applying a calibrated magnetic signal at 30 Hz.

GMR sensors are magnetic-field dependent resistors. To measure magnetic field strength, an input voltage is applied to the GMR, and the output voltage is recorded (Fig. S1). The GMR output voltage varies sigmoidally as a function of the in-plane component of the magnetic field (Fig. 1B). The sensor is configured such that very weak magnetic fields, around zero, result in outputs constrained to the steep linear part of the curve, thereby maximizing the dynamic range in the region of interest. In the linear part, the slope is 1.8%/mT, corresponding to a sensitivity of 10 to 25 Volt_out_/(Volt_in_xTesla). The noise spectrum at a typical input voltage of 0.5 V leads to sensitivities of 7 nT/√Hz at 10 Hz, 2 nT/√Hz at 100 Hz and 370 pT/√Hz in the thermal noise regime above 1 kHz (Fig. 1C).

### Experimental setup

We performed in vivo recordings in primary visual cortex of anesthetized cats (see STAR Methods). Figure 2 shows a schematic representation of the experimental setup. A magnetrode was inserted into the tissue to a depth of less than 1 mm from the cortical surface using micromanipulators under microscope inspection. The magnetrode was sensitive to fields orthogonal to the tip, that is, parallel to the cortical surface. A tungsten electrode was targeted to be less than 1 mm from the magnetrode, to simultaneously obtain an independent electric recording. Recordings were performed without shielding. To physiologically activate the recorded brain area, a flash of light was presented directly to one eye of the cat. The duration of light stimulation was either 100 ms or 500 ms, with a variable inter-stimulus interval of 0.9 to 1.5 s to avoid adaptation or entrainment. The stimulus was presented 1000 times. The output signals from the tungsten electrode and the magnetrode were preprocessed and averaged with respect to stimulus onset, to calculate the event-related potential (ERP) for the electrode and the event-related field (ERF) for the magnetrode (see STAR Methods).

**Figure 2.**
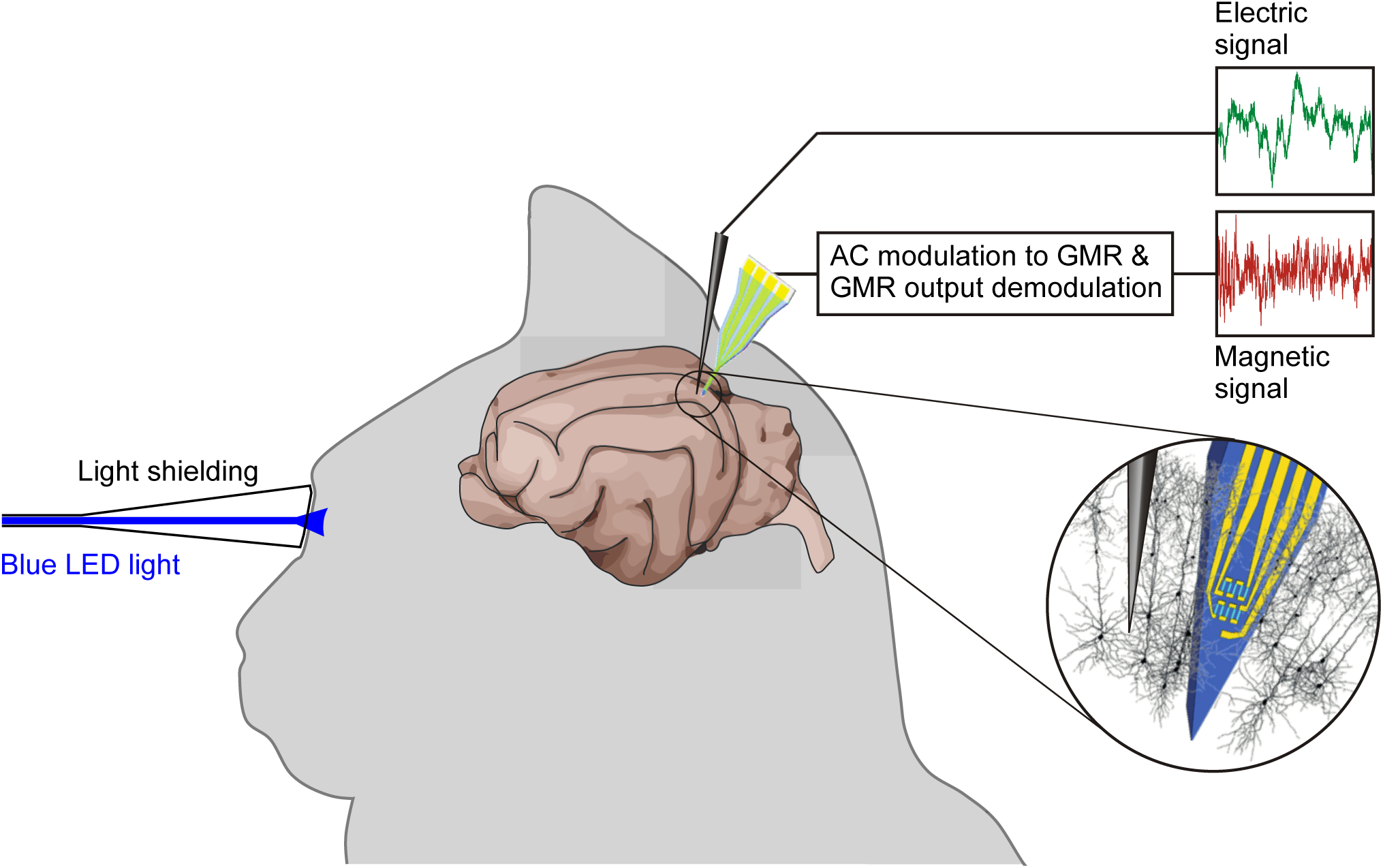
Experimental setup. Recordings were performed in primary visual cortex of the anesthetized cat. To activate the area, a visual stimulus was applied directly to the contralateral eye using blue LED light. The magnetrode, containing the GMR sensor, was positioned within visual cortex. A tungsten electrode was targeted to be less than 1 mm from the magnetrode, to simultaneously obtain an independent electric recording. The zoomed-in inset illustrates the expected configuration of the magnetrode and electrode in the neuropil. The output signal from the GMR sensor was demodulated. Subsequently, the GMR and the electrode signal were amplified, filtered and digitized.

### Estimation of the expected field strength

We estimated the field strength that we could expect, when recording magnetic fields inside the neuropil. As a starting point, we used the well-established magnetic fields recorded with MEG. When MEG signals are recorded from human subjects presented with visual stimuli, event-related fields (ERF) can be obtained with typical amplitudes in the range of 50 fT (Salmelin et al., 1994). For these MEG sensor-level field strengths, detailed models of the underlying sources estimate dipole strengths in the range of 10 nA*m (Hämäläinen et al., 1993; Murakami and Okada, 2006). We constructed a model of an ensemble of neurons, which can produce such dipole strength, to then calculate the field strengths expected for magnetrode measurements very near or inside this neuronal ensemble. We simulated a square array of 10,000 aligned neurons with a mean center-to-center separation of 5 μm. In each neuron, a current was simulated, such that the ensemble of neurons appeared as a dipole of 10nA*m, when recorded from a large distance. The corresponding difference in electric potentials was simulated to occur over a distance of 200 μm. This distance was estimated from current-source density measurements in cat visual cortex in response to visual stimuli (Mitzdorf, 1985). The currents in the neuronal ensemble gave rise to a magnetic signal of 50 fT at a distance of 6 cm, 800 fT at 1.5 cm, 126 pT at 1 mm, 1.3 nT at 100 μm and 2.3 nT inside the neuronal ensemble. Thus, these simulations predict that magnetic field measurements within or in close proximity to the activated neurons will give ERFs in the range of a few nano-Tesla.

### Separation between magnetic signal and electrical contamination

When magnetrodes are introduced into the neuropil, they might face direct capacitive coupling to electric currents flowing in the neuropil. Therefore, we developed a measurement scheme that suppressed this capacitive coupling. In this scheme, the GMR sensors were fed with alternating current (AC, Figure S2) with frequencies in the range of 20-80 kHz, and the sensor output was demodulated separately for components that were in-phase with the AC modulation and those that were out-of-phase.

The currents fed to the GMR during the AC measurement scheme are not expected to directly influence neurons in the vicinity of the magnetrode. We estimated, for a typical AC current, the resulting magnetic and electric field intensities induced in the neuropil, and they were several orders of magnitude below thresholds for neuronal stimulation (see STAR Methods).

We used two phantoms, one to generate purely magnetic fields, and another one to generate purely electric fields. When the input to the GMR was a time-varying magnetic field, the GMR output reflected this almost purely on the in-phase component (Fig. 3A). By contrast, when the input to the GMR was a time-varying electric field, the GMR output reflected this primarily at higher frequencies and then primarily in the out-of-phase component (Fig. 3B). Electric fields also induced a small in-phase component, presumably due to a mixing in the silicon substrate.

**Figure 3.**
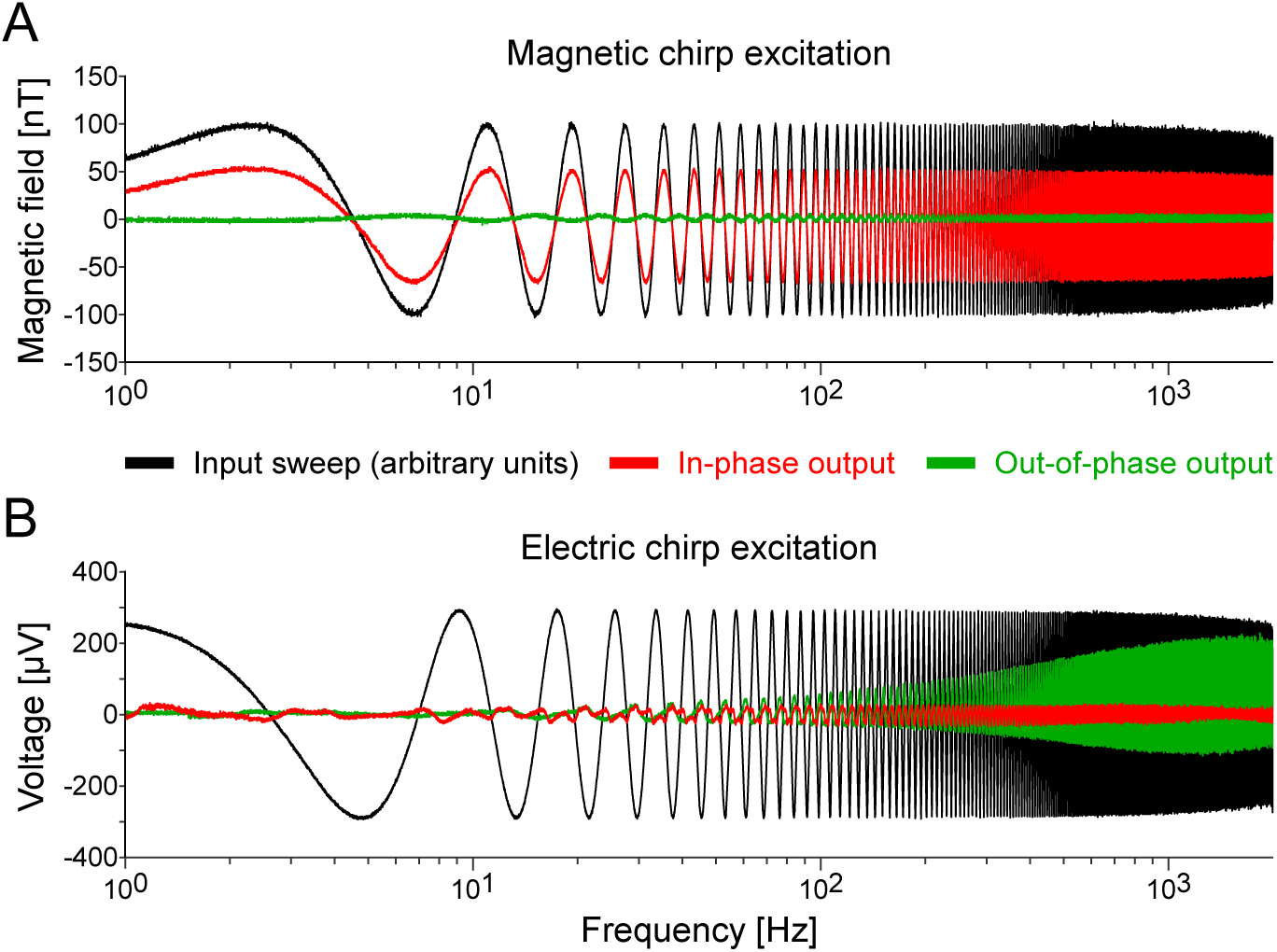
GMR output in AC mode to electric and magnetic field inputs. (A) The black line shows in arbitrary units the magnetic field input to the GMR, generated by a respective phantom. The input signal was an exponential chirp, i.e. a sinusoidal current with frequency varying exponentially from 1 Hz to 2 kHz. The GMR output was demodulated, and the in-phase output is shown in red, the out-of-phase output in green. Magnetic input is expected to be reflected primarily in the in-phase output, which is confirmed. (B) Same as (A), but with an electric field input (black line, in arbitrary units). Electric field input is expected to be reflected more in the out-of-phase output, which is confirmed, particularly for higher frequencies.

### Validation of the magnetic nature of in vivo recordings

The phantom measurements provided GMR in-phase and out-of-phase outputs for all physiologically relevant frequencies of electric or magnetic field input. Thereby, they provided a transfer function for electric fields and a transfer function for magnetic fields.

In order to estimate contamination from electric field in vivo, we used the ERP recorded in one session (cat 2B) and convolved it with the transfer function estimated for electric fields in the phantom measurements. This provided an estimate of the GMR output that would be expected if the input were purely an electric field with the waveform of an ERP (Fig. 4A). In this case, the GMR out-of-phase component (green line) was larger than the in-phase component (red line).

**Figure 4.**
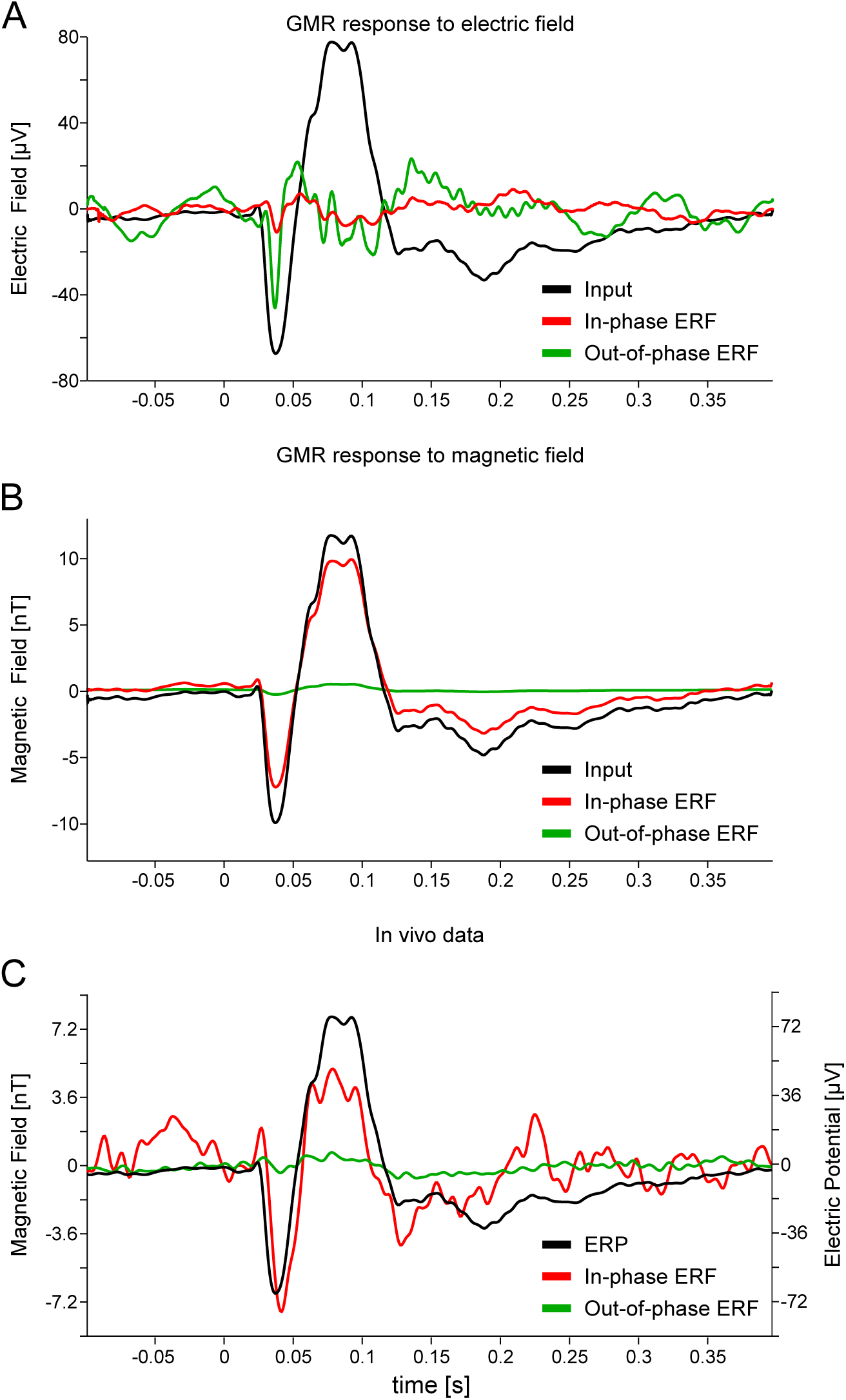
Validation of the magnetic nature of in vivo recordings. (A) GMR output after in-phase (red) and out-of-phase (green) demodulation, which would be expected (based on the phantom measurements shown in Fig. 3), if the input were a purely electric field with the waveform of an ERP (black). (B) Same as (A), if the input were a purely magnetic field with the same waveform. (C) Experimentally observed GMR output after in-phase (red) and out-of-phase (green) demodulation. The ERP recorded simultaneously from an independent tungsten electrode is shown in black. The ERP and the GMR outputs are averages over 1000 stimulus repetitions.

Subsequently, we convolved the same ERP waveform with the transfer function for magnetic fields. This provided an estimate of the GMR output that would be expected if the input were purely a magnetic field with the waveform of an ERP (Fig. 4B). In this case, the in-phase component (red line) was substantially larger than the out-of-phase component (green line). We used the ERP waveform for both simulations, to aid direct comparison and to avoid circularity, when we compare, in the next step, the simulated GMR outputs to the experimentally observed GMR output.

The observed GMR output (Fig. 4C) showed a substantially larger in-phase component (red line) than out-of-phase component (green line). This pattern corresponds to the pattern estimated for magnetic field input (Fig. 4B), which suggests that the GMR output is mainly determined by the neuronally generated magnetic fields. The magnetic field input is primarily reflected by the in-phase component of the GMR output. Therefore, in the following, we refer to the in-phase component of the GMR output as event-related fields (ERFs), and we compare them to the event-related potentials (ERPs) recorded simultaneously through the tungsten electrode.

### Comparison between simultaneously recorded event-related fields (ERFs) and event-related potentials (ERPs)

Figure 5A shows the ERF and Figure 5B the simultaneously recorded ERP for the recording in the first animal (cat 1) with a visual stimulus duration of 100 ms. Figure 5C shows a magnification of the data with the ERF (red) and ERP (green) scaled and superimposed to facilitate comparison. The ERF showed a magnetic response starting 20 ms after stimulus onset, corresponding to the conduction delay between the retina and the primary visual cortex. The ERF was characterized by a strong negative component at 36 ms and a positive peak around 61 ms. The peak-to-peak amplitude was 2.5 nT. The onset of the ERP was comparable to the magnetic one, with a trough at slightly shorter latency and a peak at similar latency as the magnetic signal. Figure 5D shows the Pearson correlation coefficient between ERF and ERP as a function of time lag, with positive lag values indicating that the ERF lagged the ERP. The correlation function peaked at a value of approximately 0.55, for a lag of approximately 2 ms. The side peaks and troughs are due to the partially rhythmic nature of the ERP and ERF.

**Figure 5.**
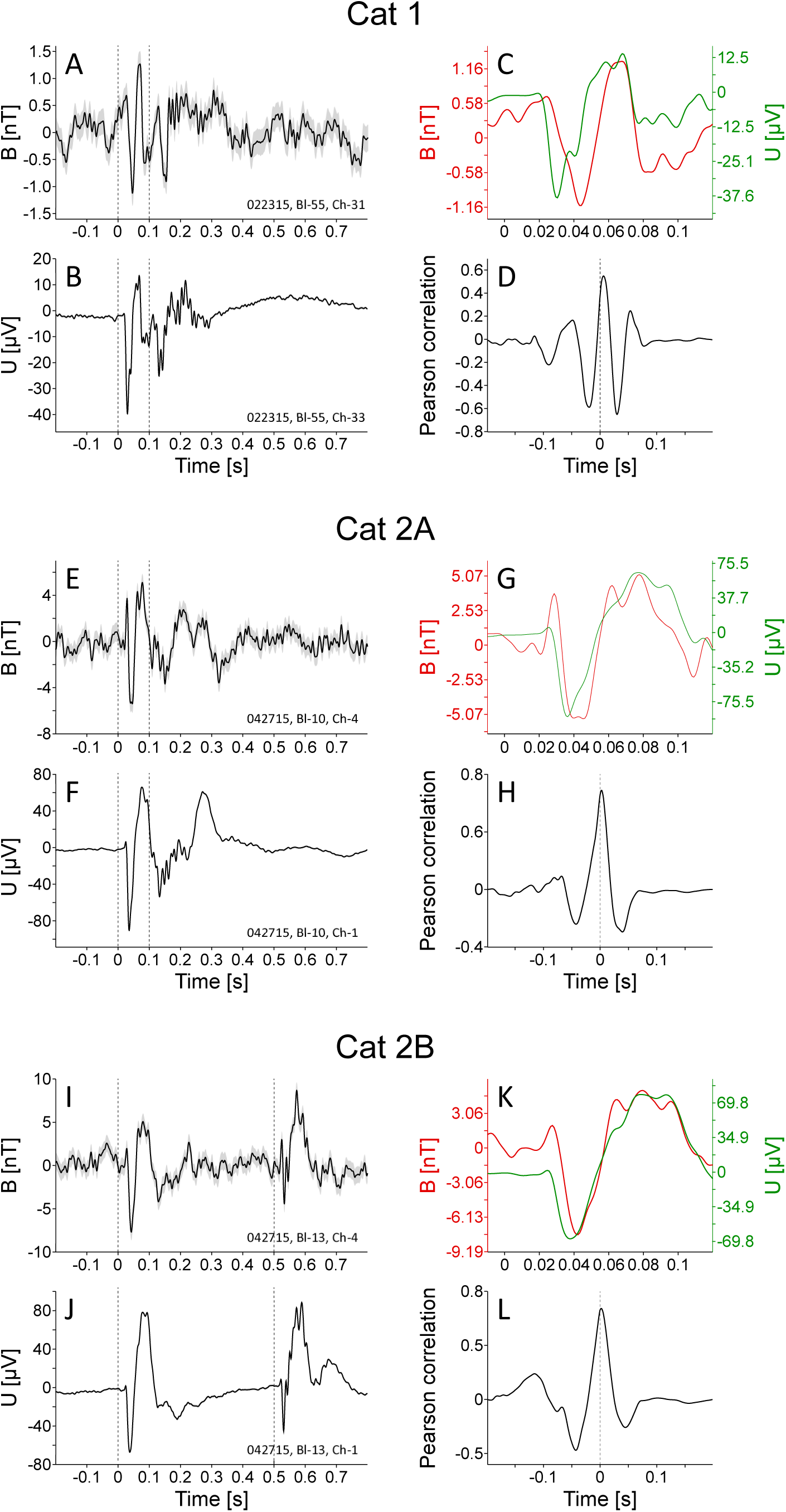
Comparison between simultaneously recorded event-related potentials (ERPs) and event-related fields (ERFs). (A) ERF obtained in cat 1 by averaging the GMR in-phase output over 1000 stimulus repetitions. The dashed vertical lines indicate onset and offset of the 100 ms long visual stimulus. (B) ERP obtained simultaneously by averaging the signal from an independent tungsten electrode, over the same 1000 stimulus repetitions. For both the ERF and the ERP, the gray shaded regions show ±1 SEM. The error region of the ERP can be visually appreciated by magnifying the figure. (C) Direct comparison of the waveforms of the ERF (red) and the ERP (green). (D) Pearson correlation coefficient between the ERF and the ERP as a function of time lag. (E-H) Same as (A-D), but for a recording session in cat 2. (I-L) Same as (E-H), but for a separate recording session in cat 2, using a 500 ms long visual stimulus.

Similar results were obtained in two separate recordings from another animal (cat 2A and cat 2B). Figure 5E-H shows the results for one recording site and a visual stimulus duration of 100 ms. Figure 5I-L presents the data from another recording site later in the experiment, with a visual stimulus duration of 500 ms. With the longer stimulus duration, the on and off responses were clearly separated, as evident in the magnetic and electric recordings. The signal amplitude of the magnetic (and of the electric) recordings was larger in cat 2, with peak-to-peak amplitudes of approximately 10 nT. Similar to cat 1, the electric signal had a shorter latency than the magnetic signal, but in cat 2 the difference was only a few milliseconds. The cross-correlation functions between the ERFs and ERPs of cat 2 showed peak values around 0.85 at a lag of 2-3 ms.

### Evaluation of signal quality

To further characterize the magnetic responses, we determined two metrics of signal quality. In a first approach, we calculated a simple metric of signal-to-noise ratio (SNR), based on the mean squared ERF or ERP (see STAR Methods for details). When this SNR was determined for ERFs based on averaging all 1000 trials, it reached values between 12 and 17 (Fig. 6A). When ERFs were based on averaging increasing numbers of trials, they reached significance at 229, 103 and 95 trials, for recording sessions cat 1, cat 2A and cat 2B, respectively (Fig. 6B; bootstrap test, see STAR Methods). For ERPs, it reached maximal values between 30 and 36 and was significant for single trials (Fig. 6C).

**Figure 6.**
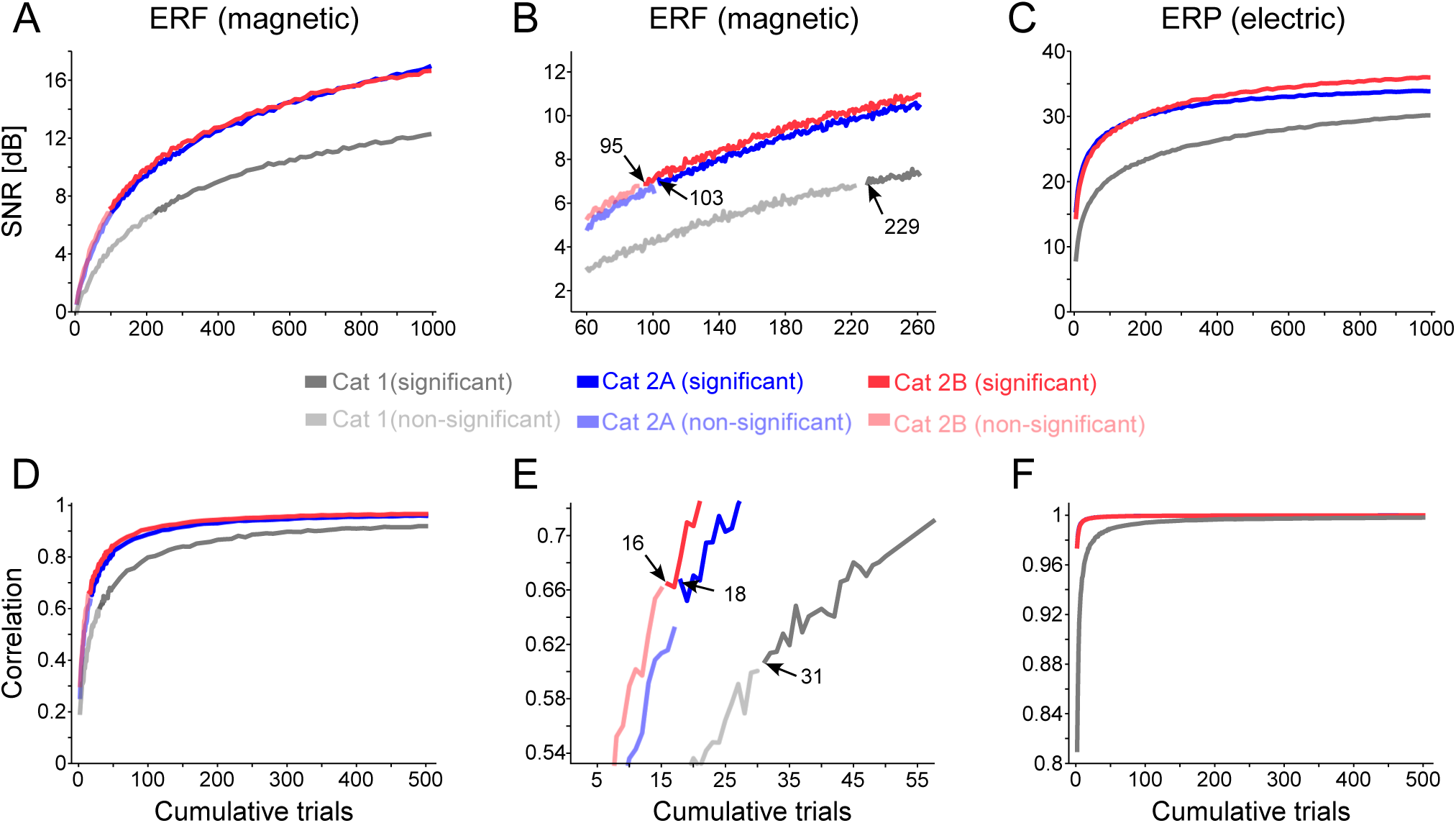
Evaluation of signal quality. (A) Signal-to-noise ratio (SNR) of the ERF as a function of the number of trials that were averaged. As specified in the color legend, different colors refer to different recording sessions, and color saturation indicates significance. (B) Same as (A), but zoomed in on the transition to significance. (C) Same as (A), but for the simultaneously recorded ERP. (D) Pearson correlation coefficient between a template ERF averaged over 500 trials and a subset-ERF averaged over the number of trials specified on the x-axis. Template and subset ERF always averaged non-overlapping sets of trials. (E) Same as (D), but enlarged around the transition to significance. (E) Same as (D), but for the simultaneously recorded ERP. Note that metrics for ERFs and ERPs are shown with different y-axis scales. The correlation values for ERPs of cat 2A and cat 2B (F) are very similar and largely overlap.

While the SNR metric is simple, it is not very sensitive. Therefore, in a second approach, we quantified how many trials had to be averaged for the ERF or ERP to assume its final shape. We first selected a random half of all trials to calculate a template shape. We then averaged increasing numbers of the remaining trials and calculated the correlation between the resulting shapes and the template shape. When the correlation was determined for ERFs based on averaging all remaining 500 trials, it reached values between 0.92 and 0.97 The correlation reached significance for 31, 18 and 16 trials, for recording sessions cat 1, cat 2A and cat 2B, respectively (Fig. 6E; bootstrap test, see STAR Methods). For ERPs, it reached maximal values of 0.99 and was significant for single trials (Fig. 6F).

## DISCUSSION

In summary, we have shown that magnetrodes based on spin electronics can be used to record in vivo magnetic signals originating from neuronal activity. This was possible, because GMR sensors combine a small size of a few tens of microns with sufficient magnetic field sensitivity. Magnetic field recordings inside the tissue offer unique opportunities, because they are reference-free, they measure the direction of magnetic fields and thereby of the underlying (intracellular) current flows, and because these magnetic fields are not smeared by intervening neuropil. In vivo magnetic field measurements might contribute to a better understanding of the commonly recorded extracranial MEG signal. There are also efforts to record neuronally generated magnetic fields by means of magnetic resonance imaging (MRI) (Bandettini et al., 2005; Körber et al., 2013), and our magnetrode recordings provide ground-truth measurements for this.

We would like to highlight the potential utility of GMR-based sensing of neuronal activity for recordings from untethered implanted devices. Implanted recording probes play an important role in many neurotechnological scenarios. Untethered probes are particularly intriguing, as they avoid connection wires and corresponding limitations (Seo et al., 2016). Yet, for untethered probes to be maximally useful, they need to be tiny, and this results in a fundamental problem for electrical recordings. Electrical recordings require two electrochemical interfaces with sufficient distance, such that the electrical potential difference does not become vanishingly small. The necessary distance restricts the size to which untethered devices based on electric recordings can be reduced. Magnetic field recordings do not suffer from this problem, because they require merely a singular GMR. Thus, magnetrode-based untethered recordings, while challenging, might provide a unique combination of recording and transmitting modalities for future neurotechnology.

We revealed visually evoked magnetic fields by averaging over multiple stimulus repetitions. This was possible, because the underlying postsynaptic potentials (PSPs) are long-lasting compared to their temporal jitter across trials. Thereby, PSPs temporally superimpose in the cross-trial average. This holds not only for PSPs of one postsynaptic neuron, but for PSPs of many neurons in the vicinity of the magnetrode. Thus, the ERF became detectable due to effective summation of the PSP-related magnetic fields across neurons and across trials. ERFs in the different recording sites reached significance after averaging 16-31 trials (Fig. 6E). Thus, moderate improvements in sensitivity and shielding will likely enable detection of ERFs on single trials. If the detection of single-trial ERFs will succeed, also the detection of magnetic fields corresponding to single action potentials (APs) appears realistic. AP amplitudes, when recorded with electrodes close to the cell body, substantially exceed ERP amplitudes. This is likely due to the fact that each AP reflects massive transmembrane currents that move the transmembrane voltage across the cell body from -60 mV to +30 mV. Whether these strong currents generate detectable magnetic fields crucially depends on their spatial symmetry and temporal simultaneity. If all involved currents flew simultaneously and with spherical symmetry, they would generate no detectable magnetic field. However, it is known that APs emerge in the axon hillock and retrogradely invade the cell body and sometimes the dendrites (McCormick et al., 2007; Stuart et al., 1997). Thus, APs are likely magnetically visible.

Single extracellular metal microelectrodes typically record spikes from merely a handful of neurons, because insulating cell membranes isolate them from the hundreds of neurons in their immediate vicinity (Buzsáki, 2004). Magnetic fields corresponding to action potentials, that is “action fields” or AFs, should travel from neurons to the magnetrode without attenuation. This might enable AF recordings from tens or even hundreds of neurons from the vicinity of the magnetrode. The separation of spikes originating from these many neurons would be a challenge. Electrophysiological recordings allow separation of a handful of spikes, based on the relatively stereotypical spike waveform of a given neuron and the fact that millisecond-precise spike coincidences of neighboring neurons occur with a very low probability of 0.01 – 0.001 (Jia et al., 2013; Kohn and Smith, 2005). Magnetic recordings would be able to benefit from the same factors, and in addition from the vectorial nature of magnetic sources and the corresponding vectorial sensitivity of the sensors. Sensors specific for the three spatial dimensions could be combined on a single magnetrode to estimate the 3D position of each neuronal source relative to the magnetrode.

Importantly, the magnetrode as presented here, without further modifications or improvements, can provide useful ERF measurements even in an unshielded environment after averaging over merely 16-31 stimulus repetitions. These values are similar to the number of 30 stimulus repetitions, which has been estimated as the minimum to obtain a consistent visually evoked ERP from human EEG recordings (Thigpen et al., 2017). Thus, for event-related experimental designs, the fundamental benefits of magnetic in vivo recordings can now be exploited. ERFs can be used to localize the underlying ionic currents, without smearing by intervening cell membranes. In fact, ERFs are most likely dominated by intracellular currents, rather than by the extracellular return currents measured as ERPs. Thereby, magnetic field recordings could greatly refine current-source density estimates, measurements of increasing importance (Lakatos et al., 2016). The vectorial nature of the ERF recordings allows the identification of the current flow direction after combination of merely three GMR sensors. Obtaining similar information from ERP recordings requires measurements with a dense 3D grid of electrodes.

Magnetrodes also allow for an elegant combination of magnetic field recordings with magnetic resonance imaging (MRI). Modern high-field MRI can provide structural images of the living brain with sub-millimeter resolution. The spatial information in MRI is based on spatial gradients in magnetic field strength and the corresponding spatial gradients in the Larmor frequency, that is, the frequency at which protons re-emit radio-frequency energy. This frequency can be easily obtained from magnetrode measurements, and will thereby permit very precise co-registration of the magnetrode with the position in the MRI that corresponds to the measured Larmor frequencies. Finally, the magnetrode can be used to perform MR spectroscopy of the immediate magnetrode surround (Guitard et al., 2016), thereby combining the recordings of rapidly changing neuronal currents with recordings of neurotransmitters and metabolites.

The in vivo magnetic field measurements presented here have a sensitivity that is still below the conventional metal microelectrode. We would like to compare this situation to the early days of MEG measurements in humans, when its sensitivity was probably far below EEG. Today, EEG is preferred, when e.g. large cohorts of subjects are measured (Dikker et al., 2017) or when combined with fMRI (Scheeringa et al., 2011); MEG is preferred, when spatial localization and measurements of deep sources is essential (Gross et al., 2002). Similarly, electrophysiological and magnetophysiological in vivo recordings will be complementary. Conventional microelectrodes and modern MEMS-based multi-contact electrodes will continue to be indispensable workhorses for neurophysiology. At the same time, magnetrodes will allow recordings that are different in nature and thereby offer distinct advantages for answering specific questions.

## AUTHOR CONTRIBUTIONS

Conceptualization, M.P.L., C.F. and P.F.; Methodology: L.C., J.T.R., J.P.A., J.V. and V.T.; Investigation: T.W., C.M.L., J.N., P.J., L.C., J.P.A., J.V., S.C., P.P.F., C.F., P.F. and M.P.L.; Analysis: T.W., V.T., J.N., C.M.L. and P.F.; Writing – Original draft: M.P.L., T.W. and P.F. Funding Acquisition: M.P.L., P.P.F. and P.F.

## ACKNOWLEDGMENTS

This work has been funded through the EU Project Magnetrodes (FP7-ICT-2011 project 600730) and through the Magsondes project by RTRA-Triangle de La Physique. This work was partly supported by the french RENATECH network. INESCMN acknowledges FCT funding through project EXCL/CTM-NAN/0441/2012 and the IN Associated Laboratory. ESI acknowledges funding through the DFG (FOR 1847, SPP 1665, FR2557/5-1-CORNET), the EU (HEALTH F2 2008 200728, HBP), the NIH (HCP WU-Minn Consortium, NIH grant 1U54MH091657), and the LOEWE program (NeFF).

## STAR Methods

### CONTACT FOR REAGENT AND RESOURCE SHARING

Myriam Pannetier-Lecoeur, SPEC, CEA, CNRS, Université Paris-Saclay, CEA Saclay, 91191 Gif-sur-Yvette Cedex, France.

### EXPERIMENTAL MODEL AND SUBJECT DETAILS

The animal experiments were approved by the responsible government office (Regierungspräsidium Darmstadt) in accordance with the German law for the protection of animals. Two adult cats (1 male, 1 female) were used for visual neuroscience experiments, after which magnetrodes were tested.

### METHOD DETAILS

#### Probe fabrication process

The GMR stack is deposited by sputtering on a commercial silicon substrate of 700 μm, insulated by a 1 μm thick SiO2 layer. The deposition is made by DC sputtering at a partial Argon pressure of 5 10^−3^ mbar. The GMR stack has the following composition (starting from the top and with thicknesses indicated in nm in parenthesis):

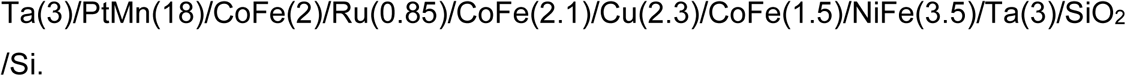

This structure contains on top the pinned layer, composed of an artificial antiferromagnet (PtMn/CoFe/Ru/CoFe).

The wafer is annealed at 300°C in a field of 1 T applied along the wafer plane and meant to set the magnetization of the PtMn/CoFe layer, which is the magnetic reference layer. After deposition and annealing, the wafer substrate is ground down to 200 μm by mechanical grinding (GDSI, USA).

The GMR stack is patterned by optical lithography. Shipley 1813 resist has been used for the entire process except the deep reactive ion etching (DRIE) process. Spin coating is realized at 5000 rpm in a clean room (which leads to a resist thickness of 1.3 μm) and soft baked at 110°C for 3 min on a hot plate to evaporate the coating solvent and harden the resist. The UV exposure process is realized on a MJB3 or MJB4 (Karl Süss, Süss Microtec, Germany) mask aligner that makes physical contact between photomask and sample. Both aligners have a wavelength of 365 nm and 5 mW/cm^2^ and 10 mW/cm^2^ power respectively. Exposure time depends on the steps (etching, contacts, passivation) and are detailed in Table 1. After exposure, the sample is rinsed in a developer to remove the exposed parts and submitted to a second bake on a hot plate. All parameters for the photolithography processes (etching of the GMR, contact deposition, passivation layer and silicon etching by DRIE) are given in Table 1. In a first step, the GMR element is etched by ion milling under Argon gas at 10^−4^ mbar, for a beam current of 7 mA and 90 W RF powered for 20 min.

Contacts are realized by a lift off process where a bilayer of Ti (15 nm)/Au (150 nm) is deposited by electron beam evaporation at 10^−8^ mbar base pressure, and 65 mA beam current for Ti and 330 mA beam current for Au.

Each GMR element is contacted on its short end (Fig. 1), along the entire height and width, with the current running in the plane of the stack.

An electrode made of Platinum (thickness 200 nm) was also deposited by evaporation at the tip of the probe (Fig. 1A), but could not be used during the in vivo recordings on AC mode, because of a saturation of the electrode amplifier by the AC current feeding the GMR.

Passivation of the structure is insured by sputtering Al_2_O_3_ (150 nm)/S_i3_N_4_ (150 nm) across the entire probe surface, except for the contact pads on the opposite side of the probe. Deposition of the passivation bilayers is done at 5.10^−3^ mbar, with 200 W RF power. After passivation, resistance leakage is quantified and only probes with infinite resistance are used.

Finally, DRIE was used to define the tip-shape of the overall probe. Deep etching uses the Bosch process (Laermer and Schilp, 1996), also known as time-multiplexed etching that alternates repeatedly between etching and passivation modes to create deep vertical penetration with a highly anisotropic profile. DRIE is a plasma etching mode with a gas mix of SF6 and O2, after a prior CHF3 etch used to remove the SiO2 layer. 400 cycles were used to etch down the silicon and define the final shape of the probe.

The probe was then mounted on a printed circuit board (PCB) by gluing its upper side and then contacted to the copper lines by wire-bonding. Wire-bonds were 50 μm thick and were protected by encapsulation in thin araldite glue.

The magnetoresistance ratio ΔR/R_0_, where ΔR is the maximum resistance variation as a function of field and R_0_ is the mean resistance, is 6.1% for this stack at room temperature.

#### Transport and noise characterization

Probes were characterized through magneto-transport and noise spectral density measurements at room temperature. For the magneto-transport experiments, a DC current of typically 1 mA was applied to the GMR element, whose output voltage was amplified and low-pass filtered at 30 Hz (Stanford Research SR560). An external magnetic field generated by two air coils mounted in Helmholtz configuration was applied along the sensitive axis of the GMR (i.e. parallel to the Pinned Layer main axis).

Noise spectral density measurements were performed in a Magnetically Shielded Room. Voltage supply was provided by two 9 V batteries in series to the GMR element and to an equivalent adjustable resistance. Both outputs were sent to the inputs of a low-noise amplifier (INA 103) in differential mode. The amplifier output was further amplified and filtered (SR 560 fed by battery). The total gain of the acquisition line was typically between 10,000 and 100,000, and the bandwidth was [0.1 Hz; 3 kHz]. An external magnetic field, in the μT range at 30 Hz, generated by an air coil, was used for calibration purposes.

The signal was Fourier transformed to obtain the noise spectral density over the chosen frequency band. The measured voltage signal together with the known signal generated by the calibrated coil resulted in the factor that was subsequently used to convert the voltage signal into an equivalent magnetic field signal.

#### Electronics schemes for GMR measurements

For general GMR characterization, the GMR was fed with a DC current, and the output voltage was measured through a low-noise amplifier and a band-pass filter from 0.1 Hz to 3 kHz.

For in vivo GMR measurements, we used a measurement scheme that suppressed capacitive coupling to the neuropil, which may occur in DC measurements (Amaral et al., 2011). In this scheme, the GMR sensors were fed with alternating current (AC, Fig. S1) with frequencies in the range of 20-80 kHz, and the sensor output was demodulated separately for components that were in-phase with the AC modulation and those that were out-of-phase. The in-phase signal is linked to the resistive part of the bridge (i.e. the GMR), and the out-of-phase signal relates to the capacitive coupling to the neuropil. Thus, the AC mode is designed such that magnetic fields are detected on the in-phase channel and electric fields on the out-of-phase channel.

Total gain of the acquisition line typically ranged from 500 to 1000. The frequency band for recordings was [0.3 Hz; 3 kHz] or [0.3 Hz; 1 kHz].

#### Phantom experiments

Phantom experiments were developed to separate between magnetic signal and potential electrical contamination. The setup contained two sources, one source generating a magnetic field and a second source generating an electric field. The magnetic field source consisted of an isolated wire immersed in saline solution, connected to a current source. The electric current source consisted of two wires with exposed ends in the bath, connected to a separate current source. A magnetrode was placed close to both sources within the saline bath.

Sweep signals ranging from 1 Hz to 3000 Hz were used to drive either the magnetic or the electric phantom separately. On the basis of those measurements, the transfer functions of the system to electric or magnetic inputs were calculated. The transfer functions allow to calculate the responses of the sensor to any given electric or magnetic field input (as long as the input is constrained to the 1-3000 Hz range).

#### Magnetic and electric fields generated by the AC current in the GMR

To evaluate the possible impact onto the neuropil, of the GMR feeding current in the AC mode, we performed an estimate of the magnetic and electric fields generated in close vicinity to the probe, to compare it with excitability thresholds as reported for Transcranial Magnetic Stimulation. The electric field induced in the medium when the GMR is supplied by an AC current *I* can be estimated by a simple model. Considering a gold contact line longer than the dimensions of interest, the magnetic field can be written as follows:
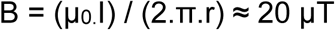

where *I* = 1 mA, and where *r* = 10 μm is the distance between the contact line and the point of interest in the bath, *μ*_0_ being the permeability of the tissues. The equation of induction links the time variations of *B* through a surface *S*, to the induced electric field E along the edge of this surface. If we consider a circular loop of radius *R* = 1 μm, then the electric field intensity is approximated by
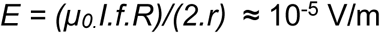

for a frequency *f* = 10^5^ Hz. This electric field intensity is several orders of magnitude below the typical values used for neuronal stimulation (100 V/m as reported in (Ilmoniemi et al., 1999)).

#### In vivo recording procedures and data analysis

Anesthesia was initiated intramuscularly with 10 mg/kg ketamine hydrochloride (Ketavet, Zoetis, Germany) and 0.05 mg/kg dexmedetomidine (Dexdormitor, Orion Pharma, Germany) supplemented with 0.04 mg/kg atropine sulfate (Atropin, B.Braun, Germany). Anesthesia was maintained after tracheotomy by artificial ventilation with a mixture of N2O/O2 (70/30%) with 0.8% isoflurane. Analgesia was maintained by intravenous infusion of sufentanil (2 μg/kg/h, Sufentanil-Hameln, Germany) together with electrolytes (3 ml/kg/h, Sterofundin, B.Braun, Germany) and glucose (24 mg/kg/h, bela-pharm, Germany). After all surgical procedures had been terminated, the animals were paralyzed by intravenous infusion of vecuronium bromide (0.25 mg/kg/h, Vecuronium-Inresa, Germany). Atropine (Atropine-POS 1%, Urspharm, Germany) was topically applied to the eye in order to dilate the pupil. Depth of anesthesia was controlled by continuously monitoring the electrocardiogram and CO2 level. Dexamethasone (Voren, Boehringer Ingelheim, Switzerland) was administered every 48 h and if needed. A craniotomy was performed around the central part of the primary visual cortex area 17 (homologue to V1 in primates, Horsley–Clarke coordinates AP - 2 to -10, ML 0 to +6). The dura mater was removed in a small window to allow easy insertion of the recording probes.

No shielding was used for the recordings, neither a mu-metal shield as in MEG, nor an aluminum shield as in a Faraday cage. This led to substantial line-noise artifacts at several frequencies (Fig. S3 and S4). Nevertheless, we reasoned that a shielding would have complicated experimentation, while its benefits would have been partly counteracted by the requirements of life support. Specifically, the animal was connected to an ECG monitor, it received intra-venous infusions, it’s body temperature was recorded via a rectal thermo probe, which was connected to a control unit, which in turn drove a heating pad. These limitations can be overcome by appropriate modifications in life support equipment or through experiments in awake animals. For the present proof of principle, averaging over multiple stimulus repetitions was effective in revealing the stimulus evoked responses even in the presence of strong noise.

Electrical recordings were performed with tungsten electrodes (1 MΩ impedance, FHC, USA). The electrode and the magnetrode were held by separate micromanipulators (David Kopf Instruments, USA) allowing for a precise positioning and careful insertion into the cortex under microscope inspection. The magnetrode was inserted first, about 1 mm below the cortical surface, and angled such that the probe penetrated the cortex as perpendicularly as possible. Subsequently, the tungsten electrode was inserted in close vicinity to the magnetrode. Given the cortical thickness of the cat, the sensors were expected to be located near cortical layer 4, the input layer. Signals from the magnetrode (after demodulation) and the electrode were recorded with a standard acquisition system (Tucker Davis Technologies, USA). To this end, signals were buffered by a unity gain headstage, high-pass filtered at 0.5 Hz, low-pass filtered at 300 Hz and digitized at 1017 Hz.

For visual stimulation, a brief (100 or 500 ms) flash of light was applied directly to the contralateral eye of the cat. This light flash (470 nm wavelength) was generated by an LED (Omicron-Laserage, Germany) and applied through a polymer optical fiber (2 mm diameter) ending close to the cornea, with an output intensity of about 2-10 mW at the end of the fiber. The fiber and the animal’s forehead were shielded with aluminum foil, to ensure that no light reached the magnetrode. This is important, because the magnetrode’s silicon substrate could be directly influenced by the light flash, i.e. by the photovoltaic effect (Mikulovic et al., 2016). However, the detected magnetic signals have a sharp deflection with a latency of 20-40 ms, which corresponds to the conduction delay from the retina to the cortex, ruling out a direct effect of the light flash on the magnetrode. To generate the light flash, the LED was controlled by the same unit that also controlled data acquisition (RZ2, Tucker Davis Technologies, USA). Several recording sessions were performed, each comprising 1000 light flash repetitions. The light flash had a duration of 100 or 500 ms depending on the session.

The inter-stimulus interval was 0.9 s plus a random time between 0 and 0.6 s to prevent adaptation or entrainment of the cortex to the repeated visual stimulus.

Offline data processing and analysis was done by custom written software and the FieldTrip toolbox (Oostenveld et al., 2011) coded in Matlab (The Mathworks, USA). Line noise artifacts at 50 and 100 Hz were removed by a second-order Butterworth bandstop filter applied in the forward direction. Additional artifacts that showed up as lines in the power spectra of the magnetic signal were removed by a DFT filter as implemented in FieldTrip. To this end, each analyzed data epoch was padded with surrounding data to a length of 10 s, a direct Fourier transform (DFT) was calculated for the specified frequencies, and the corresponding sine and cosine components were subtracted. The DFT filtering frequencies and the filtering effects are visible in Figure S4. To allow direct comparison, the same filtering routines were applied to magnetic and electric signals, even if the electric signals did not show some of the line noise components. For calculation of ERFs and ERPs, signals were low-pass filtered at 90 Hz with a 6^th^ order Butterworth filter applied in the forward direction. For the evaluation of signal quality, this was replaced by a 50 Hz Butterworth low-pass filter applied in both forward and backward direction.

Subsequently, data were segmented into trials starting 0.2 s before and ending 0.8 s after the light flash. For each trial, the mean was subtracted. Subsequently, trials were averaged to extract the stimulus-locked (i.e. evoked) brain activity. For electric recordings, these averages are referred to as event-related potentials (ERPs); for magnetic recordings, they are referred to as event-related fields (ERFs). To address both response types, we refer to event-related responses (ERRs). For the evaluation of signal quality, trials were further segmented into epochs containing the ERRs. We choose an analysis window starting 20 ms after stimulus onset and having a width of 100 ms to capture most of the response energy. To quantify the noise, for each trial, an equally long window was chosen at a random time between -500 to 0 ms relative to stimulus onset. For each epoch, a linear de-trending was performed by fitting and subtracting a linear regression. Signal processing was identical for post-stimulus (signal) and pre-stimulus (noise) epochs.

## QUANTIFICATION AND STATISTICAL ANALYSIS

### Signal quality estimation

To assess the quality of ERRs, we used two approaches, namely a metric of signal-to-noise ratio (SNR) and a correlation to a template response.

In the SNR approach, we defined:

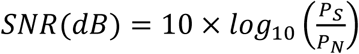

where P_s_ is the power of the ERR and PN the power of the noise. The power was quantified as the mean squared ERR in the specified epochs. Note that this definition of stimulus and noise is different from previous studies, which assume a model of additive (Gaussian) noise on top of a constant stimulus (Turetsky et al., 1988). SNR is then calculated from an estimation of the signal and noise components of the recorded stimulus evoked signal. However, we think that using the ongoing brain activity (baseline) as a measure of ‘noise’ is more intuitive because the SNR then quantifies the amount of stimulus locked activity, without making assumptions about the nature of different sources of noise. For simplicity, we keep the nomenclature of ‘signal’ and ‘noise’ for ‘stimulus evoked’ and ‘baseline’ activity, respectively.

We investigated how many trials had to be averaged to obtain a signal power, i.e. ERR power, that significantly exceeded the noise power. Starting with a single trial, we averaged increasing numbers of trials. For each number of trials, we performed the following procedure 1000 times: We randomly selected a respective subset from all 1000 available trials, and then calculated a bootstrap estimate of the 95% confidence interval (CI) of its SNR (based on 1000 bootstrap resamples of this subset). Subsequently, the upper and lower limits of the CIs from the 1000 subsamples were averaged, and the observed SNR was considered significant, if its lower average CI was larger than zero.

In a second approach, we calculated the correlation between, on the one hand the ERR obtained from a subset of trials, and on the other hand a template ERR. The template ERR was the average over 500 randomly selected trials. We tested how many of the remaining trials had to be averaged to obtain a subset-ERR that was significantly correlated to the template-ERR. Starting with a single trial, we averaged increasing numbers of trials. For each number of trials, we performed the following procedure 1000 times: We randomly selected a respective subset from the remaining 500 trials (excluding the ones used for the template), averaged them to obtain the subset-ERR, and then calculated the bootstrap estimate of the 95% CI of its Pearson correlation with the template-ERR (based on 1000 bootstrap resamples of this subset). Subsequently, the upper and lower limits of the CIs from the 1000 subsamples were averaged, and the observed correlation was considered significant, if its lower average CI was larger than zero.

## KEY RESOURCES TABLE

| REAGENT or RESOURCE | SOURCE | IDENTIFIER |
| --- | --- | --- |
| Chemicals, Peptides, and Recombinant Proteins |  |  |
| Developer AZ400K | AZ electronic materials | Cat#AZ400k |
| Developer MF 319 | Microposit | Cat#021079 |
| Primer HMDS | Microposit | N/A |
| Photoresist AZ4562 | AZ electronic materials | Cat#AZ4562 250ml |
| Photoresist Shipley 1813 | Microposit | Cat#021838 |
| Ketamine hydrochloride | Zoetis | N/A |
| Dexmedetomidine | Orion Pharma | N/A |
| Atropine sulfate | B. Braun | N/A |
| Isoflurane | CP-Pharma | Cat#1214 |
| Sufentanil | Hameln pharma plus | N/A |
| Vecuronium bromide | Inresa | N/A |
| Dexamethasone | Boehringer Ingelheim | N/A |
| Experimental Models: Organisms/Strains |  |  |
| Domestic cat (*Felis catus*) | Own breeding facility | N/A |
| Software and Algorithms |  |  |
| MATLAB | MathWorks | https://www.mathworks.com/ |
| FieldTrip toolbox | Oostenveld et al., 2011 | http://www.fieldtriptoolbox.org/ |
| Other |  |  |
| Tungsten microelectrodes | FHC | Cat#UEWMEGSECN3M |
| RZ2 BioAmp Processor | Tucker Davis Technologies | Cat#RZ2-8 |
| PZ2 Preamplifier | Tucker Davis Technologies | Cat#PZ2-256 |
| LED module (470 nm) | Omicron-Laserage | Cat# LEDMOD470.400.OEM |
| Polymer optical fiber (2 mm diameter) | Omicron-Laserage | Cat#LEDMOD.FASY.OEM |

## SUPPLEMENTAL INFORMATION

**Table S1.** Photolithography parameters.

|  | GMR | Contacts | Passivation | Deep RIE |
| --- | --- | --- | --- | --- |
| Photoresist | Shipley 1813 |  |  | HDMS + AZ4562 |
| Developer | MF319 |  |  | AZ400K+H<sub>2</sub>O (1 :4) |
| Spin coating | 5s at 500 rpm and 60s at 5000 rpm |  |  | 30s at 2000 rpm |
| Exposure | 25s at 10W/cm<sup>2</sup> |  |  | 60s |
| Development time | 60s |  |  | 240s |
| Bake 1 (before UV exposure) | 3 min at 110°C |  |  | 60 min at 90°C |
| Bake 2 (after UV exposure) | 3 min at 110°C |  |  |  |

**Figure S1.**
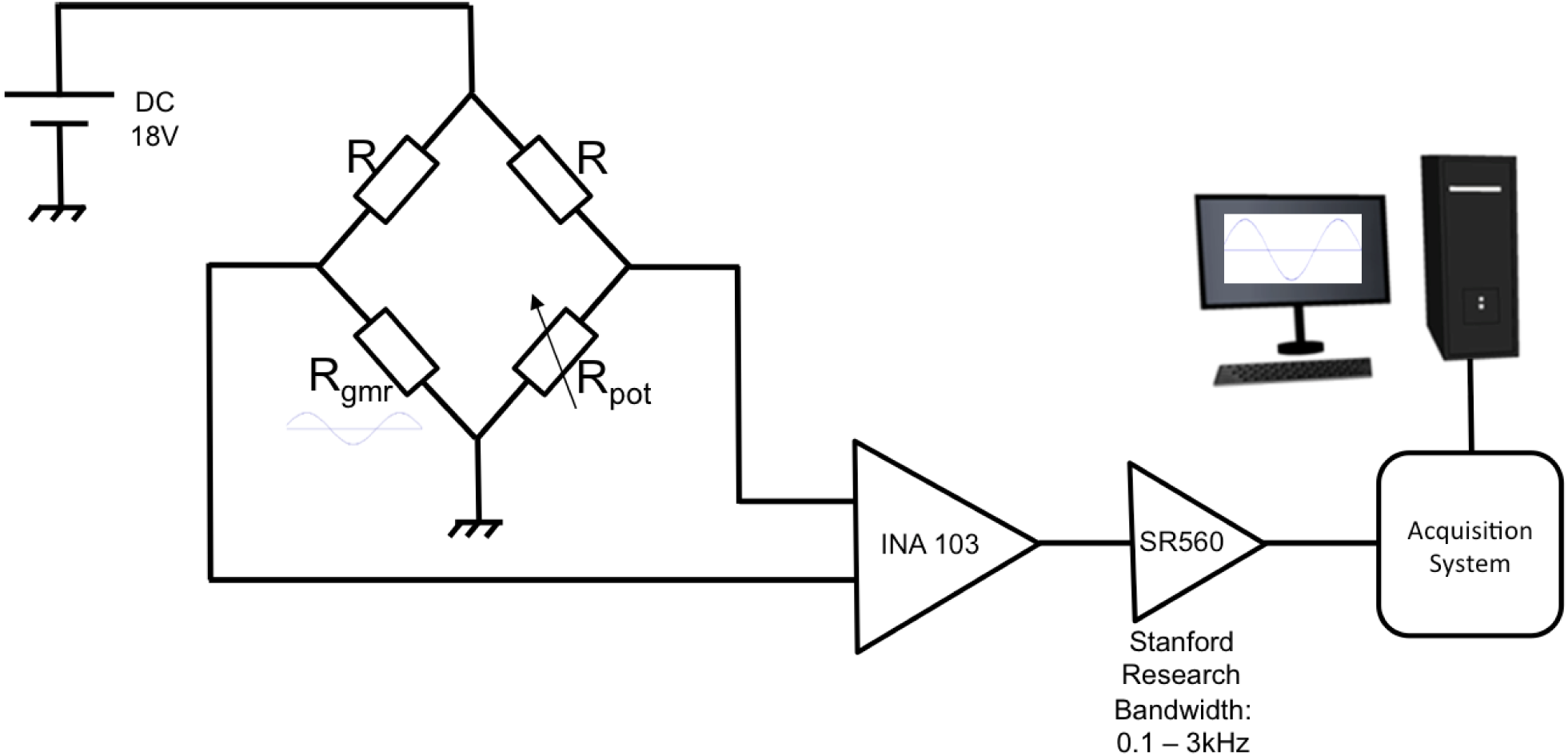
Schematic representation of DC acquisition mode. The sensor, having a resistance R_gmr_ is mounted in a Wheatstone bridge, with two identical resistances (R) of 500 Ω; a variable resistance R_pot_ is used to balance the bridge. The output of the bridge is sent through a low noise amplifier (INA 103) to a filter-amplifier (SR 560). The output voltage is collected on an acquisition card.

**Figure S2.**
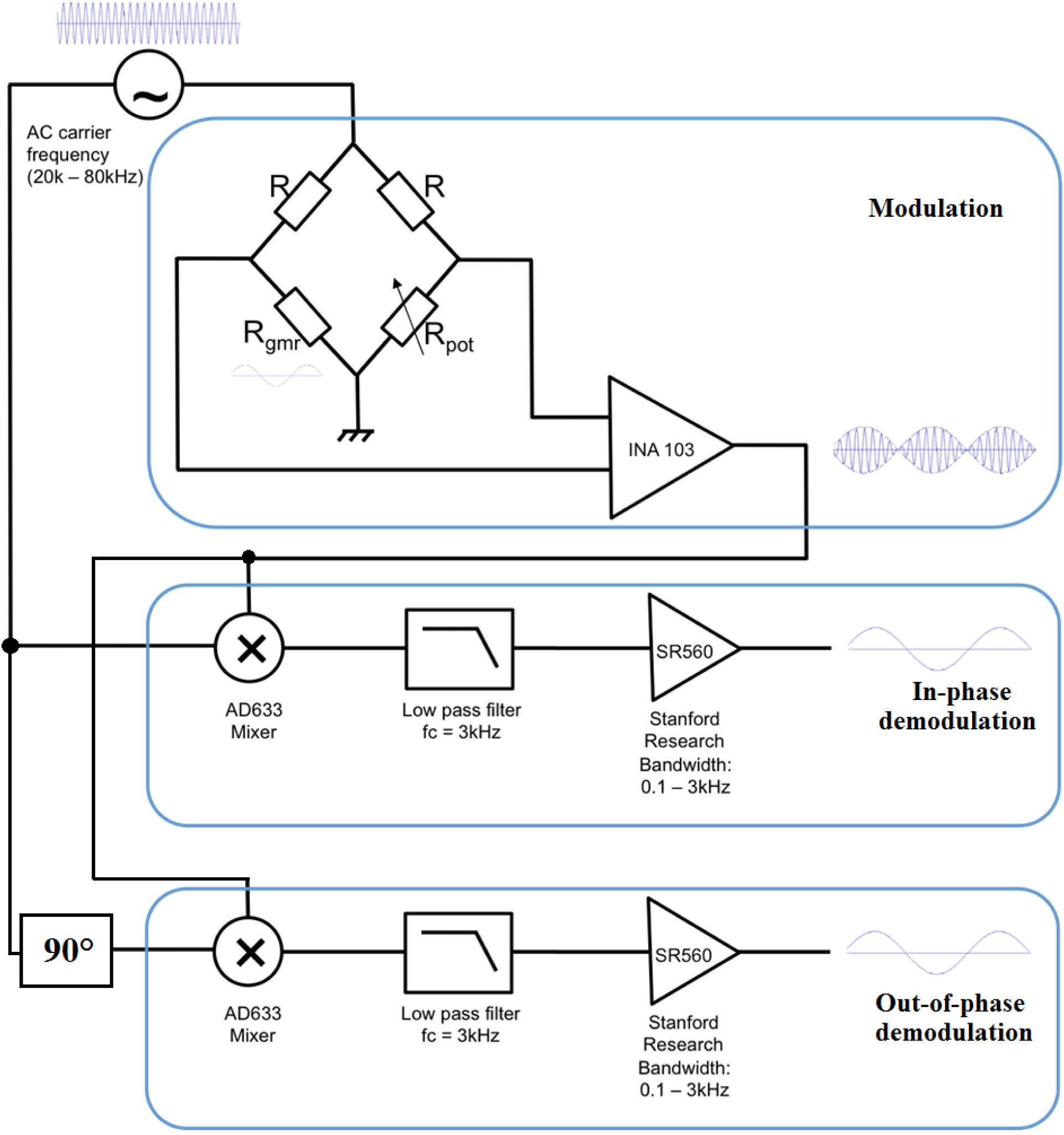
Schematic representation of AC acquisition mode. The GMR is fed with sinusoidal current of a typical carrier frequency between 20 and 80 kHz. The GMR output is a modified version of this feeding current, after amplitude modulation by the time-varying magnetic field that impinges onto the GMR. The GMR output is pre-amplified (INA 103) and demodulated by multiplication (AD633 mixers) with two carrier frequency signals, one with 0° and another with 90° dephasing. The resulting signals are low pass filtered (typically at 3 kHz) to eliminate the carrier signal contamination. Both demodulated signals, called “in-phase” and “out-of-phase”, are collected. A pure resistance change of the bridge resistance, due to magnetic field dependent changes in GMR resistance, gives an in-phase signal, whereas a capacitive change of the bridge, induced e.g. through a coupling to silicon or to the bath, induces an out-of-phase signal.

**Figure S3.**
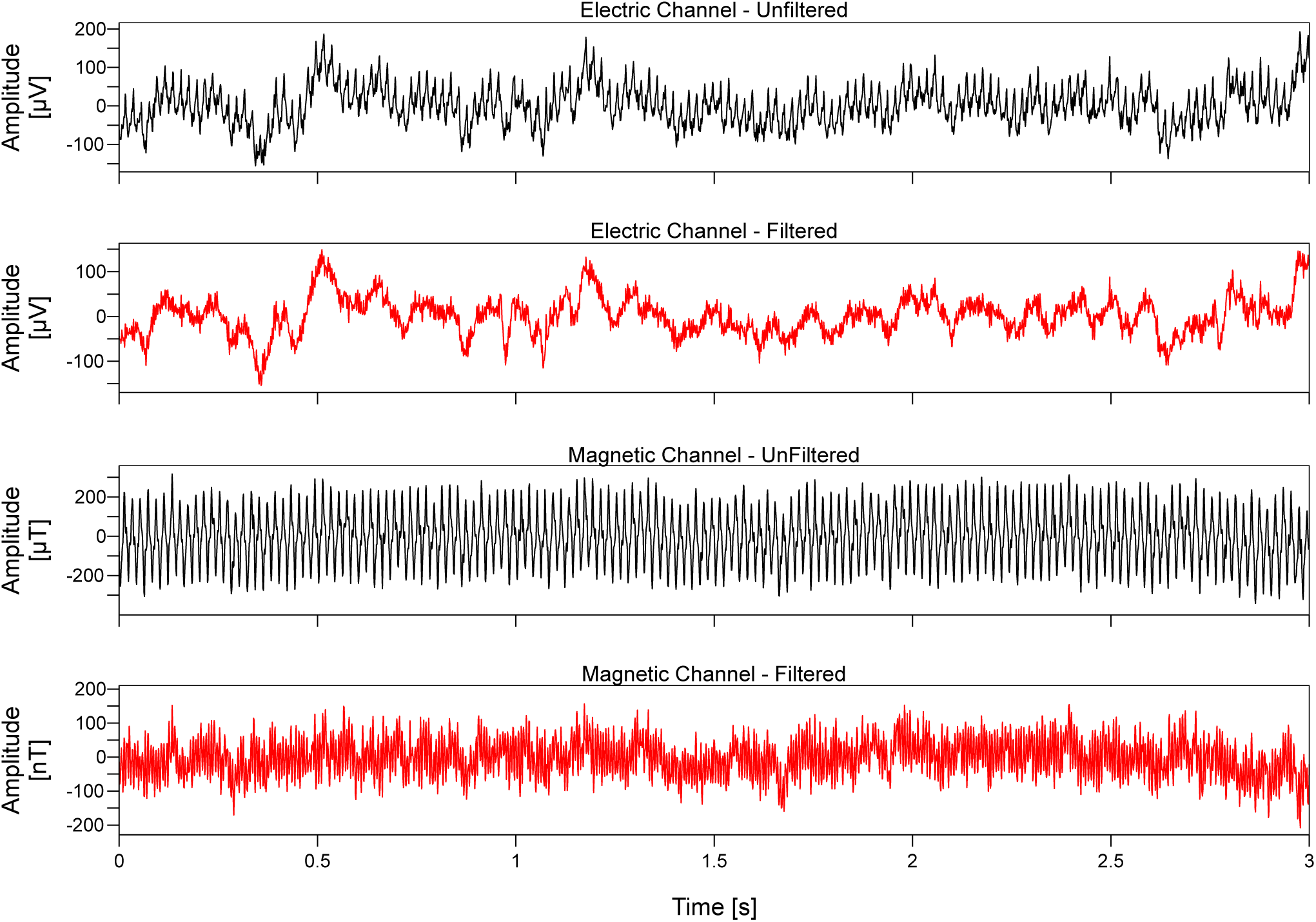
Example recordings of electric and magnetic recordings. Example traces of simultaneously recorded electric and magnetic signals, with and without filtering, as specific in the panel titles. Details of the filtering are specified in the STAR Methods section.

**Figure S4.**
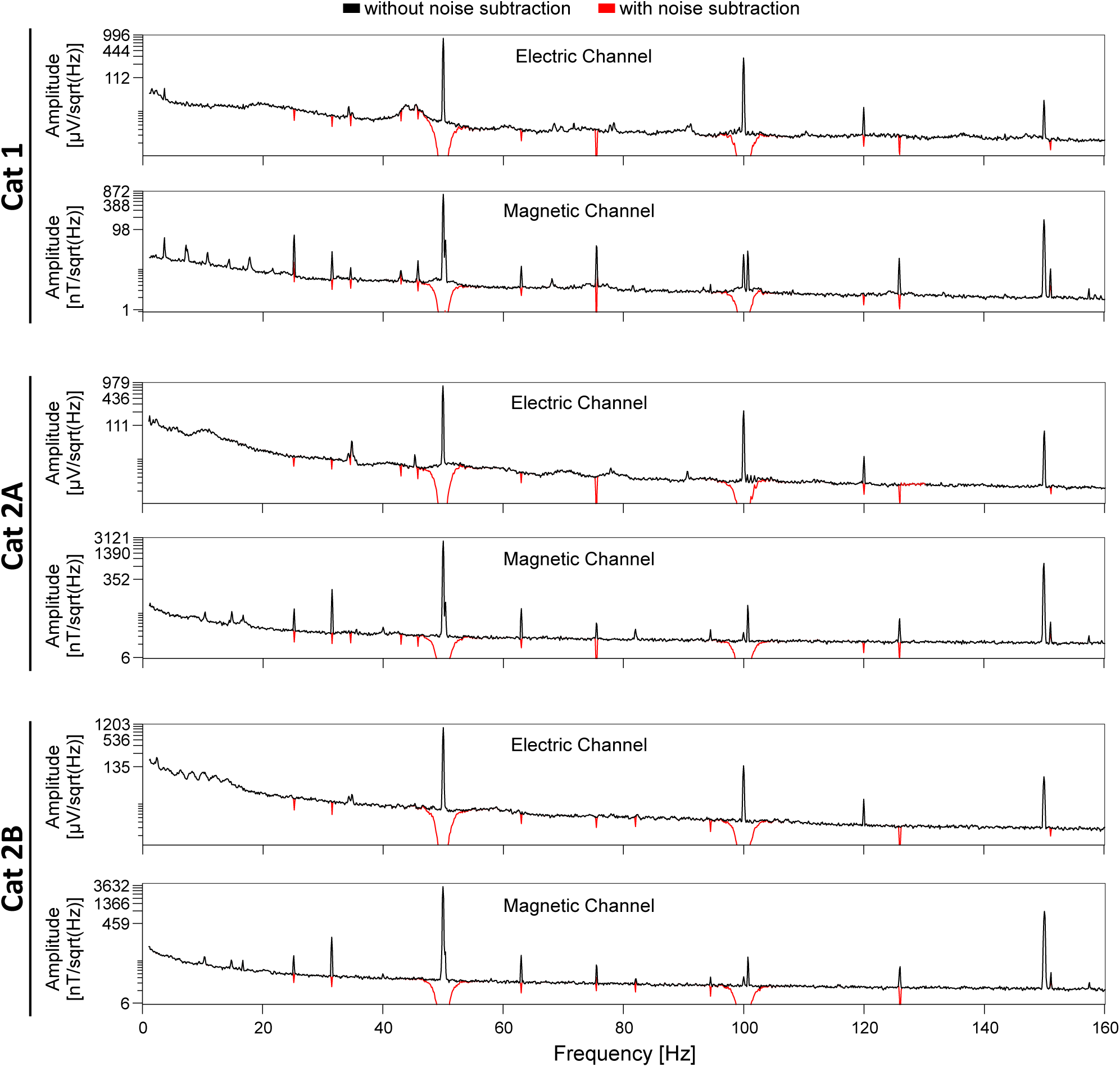
Average power spectra of electric and magnetic recordings. Average power spectra of simultaneously recorded electric and magnetic signals for the three recording sessions indicated on the left. Each panel shows in black the spectrum without noise subtraction and in red the spectrum after noise subtraction. Details of the noise subtraction are specified in the STAR Methods section.

